# Light dependent activation of the GCN2 kinase under cold and salt stress is mediated by the photosynthetic status of the chloroplast

**DOI:** 10.1101/849257

**Authors:** Ansul Lokdarshi, Philip W. Morgan, Michelle Franks, Zoe Emert, Catherine Emanuel, Albrecht G von Arnim

## Abstract

Regulation of cytosolic mRNA translation is a key node for rapid adaptation to environmental stress conditions. In yeast and animals, phosphorylation of the α-subunit of eukaryotic translation initiation factor eIF2 is the most thoroughly characterized event in regulating global translation under stress. In plants, the GCN2 kinase (*General Control Non-derepressible-2*) is the only known kinase for eIF2α. GCN2 is activated under a variety of stresses including reactive oxygen species. Here we provide new evidence that the GCN2 kinase in Arabidopsis is also activated rapidly and in a light dependent manner by cold and salt treatments. These treatments alone did not repress global mRNA ribosome loading in a major way. The activation of GCN2 was attenuated by inhibitors of photosynthesis and antioxidants, suggesting that it is gated by the redox poise or the reactive oxygen status of the chloroplast. In keeping with these results, *gcn2* mutant seedlings were more sensitive than wild type to both cold and salt in a root elongation assay. These data suggest that cold and salt stress may both affect the status of the cytosolic translation apparatus via the conserved GCN2-eIF2α module. The potential role of the GCN2 kinase pathway in the global repression of translation under abiotic stress will be discussed.

## INTRODUCTION

The translation of mRNAs by cytosolic ribosomes into new proteins is dynamically regulated by abiotic environmental conditions such as temperature (Matsuura et al., 2010; Juntawong et al., 2013;Yanguez et al., 2013), oxygen (Branco-Price et al., 2008) and light (Juntawong and Bailey-Serres, 2012;Liu et al., 2012;Missra et al., 2015;Merchante et al., 2017). Both early and more recent studies have highlighted that redox poise and reactive oxygen species can also play important roles in regulating mRNA translation in global and mRNA sequence-specific ways (Tang et al., 2003;Branco-Price et al., 2008;Khandal et al., 2009;Benina et al., 2015). The mechanisms that regulate and coordinate mRNA ribosome loading across the plant transcriptome are generally only partially understood. Of the several mechanisms regulating global translation, phosphorylation of the α-subunit of the heterotrimeric eukaryotic initiation factor 2 (eIF2) is one of the best characterized translational control events in yeast and animals (Dever et al., 1992; Donnelly et al., 2013;Hinnebusch et al., 2016). In the unphosphorylated form, eIF2 bound to GTP delivers the initiator methionyl-tRNA to the ribosomal small subunit (40S) to initiate mRNA translation (Hinnebusch et al., 2016). Upon phosphorylation by one of several kinases, eIF2α becomes a poisoned substrate of the guanine nucleotide exchange factor for eIF2, named eIF2B (Kashiwagi et al., 2018) causing global translational repression, although some mRNAs escape the global repression by virtue of specific mRNA sequence elements (Harding et al., 2000;Liu and Qian, 2014).

General Control Non-derepressible 2 (GCN2) is the only known kinase in plants that phosphorylates eIF2α (Zhang et al., 2008;Lageix et al., 2008). In the well-studied vertebrate and yeast models, the GCN2 kinase can be activated by uncharged tRNA as a consequence of amino acid starvation (Wek et al., 1995;Dong et al., 2000;Anda et al., 2017). In plants the genetic elements of the GCN2 pathway appear to be substantially conserved, although not all biochemical details have been confirmed, and few of the biochemical steps have been investigated thoroughly. Specifically, GCN2 is encoded by a single gene in Arabidopsis that functionally complements a yeast *gcn2* mutant (Zhang et al., 2003) and can be activated by uncharged tRNA *in vitro* (Li et al., 2013). Accordingly, *in planta*, the kinase is activated by inhibitors of amino acid biosynthesis such as the herbicides chlorosulfuron, glyphosate and glufosinate (Lageix et al., 2008;Zhang et al., 2008;Zhao et al., 2018), and the activation of GCN2 by herbicides can be suppressed by supplementation with amino acids (Zhang et al., 2008).

Aside from inhibitors of amino acid biosynthesis plant GCN2 kinase is activated by numerous other agents, including ultraviolet light, wounding, the ethylene precursor 1-aminocyclopropane carboxylic acid, the endogenous defense signals salicylic acid and methyl-jasmonate and bacterial infection (Lageix et al., 2008;Liu et al., 2019). What remains unclear is the nature of the biochemical signal that activates GCN2 under this variety of abiotic and biotic stresses. We recently described that GCN2 is activated by light-dependent reactive oxygen species (ROS) from the chloroplast. Even the stimulation of GCN2 by inhibitors of amino acid biosynthesis requires light and does not occur in darkness, suggesting that ROS are an essential requirement for GCN2 activation (Lokdarshi et al., 2019). A second conundrum surrounding plant GCN2 is that *gcn2* mutants have rather mild phenotypes under favorable lab conditions (Zhang et al., 2008;Lageix et al., 2008;Liu et al., 2015b) and a near-normal transcriptome (Faus et al., 2015; Lokdarshi et al., 2019). Moreover, among the various treatments that activate eIF2α phosphorylation, the herbicide chlorosulfuron is the only one that also results in a GCN2-dependent global translational repression (Lageix et al., 2008;Lokdarshi et al., 2019). In fact, the conditions that trigger eIF2α phosphorylation by the GCN2 kinase are not well correlated with the conditions under which *gcn2* mutant plants display maladaptive phenotypes.

Here we describe that the GCN2 kinase is activated by cold and salt stress in a light dependent manner. The activation of GCN2 by cold and salt can be suppressed by manipulating the status of the photosynthetic apparatus, suggesting that a chloroplastic signal contributes to the activation of GCN2. We also provide more evidence that eIF2α phosphorylation by different stresses does not always result in the same decline in polyribosome loading. However, *gcn2* mutant seedlings from two different ecotypes of Arabidopsis show reduced primary root growth under cold and salt stress, in keeping with a physiological role for the GCN2 kinase to adapt to these conditions. Taken together, these data suggest that the retrograde signaling from chloroplast to cytosol that targets protein synthesis may operate via the GCN2 kinase under cold and salt stress.

## MATERIALS AND METHODS

### Plant materials and growth conditions

*Arabidopsis thaliana* ecotype Landsberg (Ler-0), Columbia (Col-0), and homozygous *gcn2-1* mutants of the GT8359 gene trap line (Zhang et al., 2008), and homozygous *gcn2-2* mutant seeds (Lokdarshi et al., 2019) were sterilized and stratified at 4°C for 2 days. Seeds were germinated on half-strength Murashige-Skoog (1/2X MS) plant media (MP Biomedicals, cat # 2633024) with 0.65% Phytoagar (Bioworld, cat # 40100072-2) and grown under a long-day period of 16 h light (80±10 μEin m^−2^ s^−1^)/8 h dark at 22 °C and 50% humidity. Unless stated, no sucrose was added to the medium.

### Stress treatments and phenotype characterization

For cold stress treatment in dark and light, plates with 14-days-old horizontally grown seedlings were acclimated in the dark for 24 h starting at Zeitgeber time 2 (ZT2), after which they were shifted to 4 °C in the dark or light for the desired times. Dark-treated seedlings were harvested under green safe light. For salt stress treatment in the dark, plates with 9-days-old vertically grown seedlings were acclimated in darkness for 24 h starting at ZT2, after which seedlings were transferred to high salt or mock 1/2X MS salt media under green safe light and sampling was performed at the desired times. For salt stress treatment under light, seedlings were transferred to high salt (150mM NaCl), or control conditions (0.1% sucrose), or control conditions with equivalent osmolarity (300mM mannitol) starting at ZT2.

For chemical treatments with DCMU (Thermo-Fisher, cat# D2425) and DBMIB (Thermo-Fisher, cat# 271993) seedlings were sprayed with the desired amount of reagent and mock control (DMSO or water) under green safe light 30 minutes before the end of 24 hr dark acclimation. For antioxidant treatment, seedlings were germinated and grown for 10 days on 1/2X MS medium containing 0.5mM ascorbate and 0.5mM reduced glutathione.

For phenotype characterization under cold stress, 3-days-old vertically grown seedlings on 0.1% sucrose were transferred to media without sucrose and shifted to 4 °C for 30 days. For salt stress, 3-days-old vertically grown seedlings without sucrose were transferred to media with 0.1% sucrose, or supplemented with 300mM mannitol, or 150mM NaCl. Photographs were taken with a digital camera (Canon) and primary root length was measured using ImageJ (ver. 1.41; http://rsb.info.nih.gov/ij/index.html). Fresh weight measurements were performed by weighing seedlings per plate at the end of the stress treatment. Percent survival analysis for salt stress was performed by counting seedlings that showed bleached chlorophyll and no primary root growth from day 6 to day 9. All statistical analysis was performed using GraphPad Prism (ver. 8.1.2; GraphPad Software, Inc).

### Protein extraction and immunoblot analysis

Sampling for total protein extraction was done by flash freezing 14-days-old seedlings in liquid nitrogen. Seedlings were ground using a plastic pestle in a 1.5 mL tube with extraction buffer containing 25 mM Tris-HCl (pH 7.5), 75 mM NaCl, 5% (v/v) glycerol, 0.05% (v/v) Nonidet P-40, 0.5 mM EDTA, 0.5 mM EGTA, 2 mM DTT, 2% (w/v) insoluble PVP (Sigma P-6755), supplemented with 1X protease and phosphatase inhibitor cocktail (Thermo-Fisher; cat# PIA32959). Total protein content was quantified by Bradford assay (Thermo-Fisher, cat# 23236).

For eIF2α phospho-immunoblot analysis, 50 μg of total protein was separated on a 12% (w/v) SDS-PAGE gel and electroblotted onto polyvinylidene fluoride (PVDF) membrane. After 1 h of blocking at 22 °C with TBST buffer (1X Tris-buffered saline [pH7.6], 0.1% Tween-20) with 10% non-fat dry milk and 0.2% BSA, the membrane was incubated overnight at 4 °C with rabbit polyclonal phospho-eIF2α antibody (Cell Signaling, cat # 9712S) diluted to 1:5000 in 1X TBST with 0.5% BSA. Following washing with 1X TBST, 10 min each for three repeats, the membrane was incubated with horseradish peroxidase conjugated anti-rabbit IgG (Vector labs, cat# PI-1000) diluted to 1:2000 in 1X TBST with 1% non-fat dry milk for 1 h at room temperature. After washing with 1X TBST, 10 min each for six repeats, horseradish peroxidase was detected using chemiluminescence (WesternBright Quantum, Advansta) as per manufacturer’s protocol. For immunoblot with rabbit polyclonal eIF2α antibody (a gift from Dr. Karen Browning, University of Texas, Austin), 5 μg of total protein was resolved by SDS-PAGE and electroblotted onto a polyvinylidene difluoride (PVDF) membrane. Blocking and incubation with antibodies was performed as described (Dennis et al., 2009) followed by chemiluminescent detection (Lokdarshi et al., 2016). Signal intensity on immunoblots was quantified with ImageJ (ver. 1.41; http://rsb.info.nih.gov/ij/index.html).

### Polysome profiling and protein fractionation

Tissue for polysome profiling was harvested as described for total protein extraction. For polysome profiling with cold stress tissue, seedlings were ground in liquid N_2_ and 0.5g of tissue powder was resuspended in 1 mL of polysome extraction buffer (200 mM Tris-HCl pH 8.4, 50 mM KCl, 25 mM MgCl_2_, 1% deoxycholic acid, 2% polyoxyethylene 10 tridecyl ether, 50 μg/mL cycloheximide and 40U/mL RNase inhibitor (Promega Cat# N2115)) and centrifuged at 13,000 × *g* for 5 min at 4 °C. One mL of the supernatant was layered onto a 10 mL 15-50% linear gradient prepared using a Hoefer gradient maker and centrifuged at 35,000 rpm (Beckmann SW 41 Ti) for 3.5 hr at 4 °C. Absorbance at 254 nm was recorded using an ISCO UA 5 absorbance/fluorescence monitor and individual data points were extracted using the DATA acquisition software (DATAQ instruments). Polysome-to-monosome (P/M) ratios were calculated as described (Enganti et al., 2018). For polysome profiling with salt stressed tissue, 150 mg of tissue powder was resuspended in 100μl of polysome extraction buffer and centrifuged at 13,000 rpm for 5 minutes at 4 °C. 100μl supernatant was layered on a 2 ml 15-50% linear gradient prepared as above and centrifuged at 50,000 rpm (Beckmann TLS55 rotor) for 1hr 10 minutes at 4 °C. Absorbance was measured as described above.

### Hydrogen peroxide quantification

H_2_O_2_ content in seedlings was measured using the Amplex Red kit (Thermo-Fisher, cat# A22188). Briefly, 30 mg of 2-week-old seedlings were flash frozen in liquid N_2_ and ground with a plastic pestle to a homogeneous powder. Pulverized tissue was resuspended in 100 μl of sterile 1X phosphate buffered saline (PBS) and centrifuged at 17000 × g at 4 °C for 2 minutes and the supernatant was used for H_2_O_2_ measurements as per manufacturer’s protocol. Relative fluorescence was measured on a POLARstar OPTIMA plate reader (BMG LABTECH) with an excitation filter at 535 nm and emission filter at 600 nm.

### Photosynthetic efficiency measurement

The maximum quantum yield of photosystem II [Qymax= F_v_ /F_m_] was measured on a FluorCam 800MF (Photon Systems Instruments) as per manufacturer’s instructions and modifications from (Murchie and Lawson, 2013). Briefly, plants were dark adapted for 2 min (F_0_) prior to applying a saturating pulse of 1800 μEin m^−2^ s^−1^ for 0.8 sec (F_m_). Variable fluorescence (F_v_) was calculated as the difference between F_o_ and F_m_ to get the maximum quantum yield [F_v_/F_m_]. For measurements under cold stress, pots with rosette stage wild-type and *gcn2* mutant plants on soil were shifted to cold (4 °C) or left at 22 °C (mock) and measurements were taken for the indicated times. Recovery from cold was done by moving the pot back to 22 °C. For F_v_/F_m_ under salt stress, 3-days-old seedlings grown on 0.1% sucrose were shifted to 1/2X MS plant media supplemented with 150mM NaCl or no salt as control (Mock) and F_v_/F_m_ measurements were recorded as discussed above.

## RESULTS

### GCN2 kinase activation under cold stress is light dependent

Previous reports (Lageix et al., 2008;Wang et al., 2017) showed eIF2α phosphorylation as a read out of GCN2 activity under cold stress. Given that the response to cold stress is closely linked to photosynthesis (Crosatti et al., 2013;Adam and Murthy, 2014) we tested whether the activation of GCN2 under cold stress was light-dependent. In wild-type Arabidopsis seedlings subjected to 4 °C cold in the light, phosphorylation of eIF2α increased gradually and remained high for at least 2 hours of cold treatment. As expected, eIF2α phosphorylation was mediated by GCN2 (Fig. 1A). In contrast, if the cold treatment was performed in dark-adapted plants, eIF2α remained unphosphorylated (Fig. 1C). Under regular temperature conditions in the light, eIF2α-P remained steady between ZT2 and ZT4 (Fig. 1B). Additionally, under all the test conditions the overall amount of eIF2α remained unchanged (Fig. 1A-C). These results show that GCN2-dependent eIF2α phosphorylation under cold stress is light dependent.

**Figure 1.**
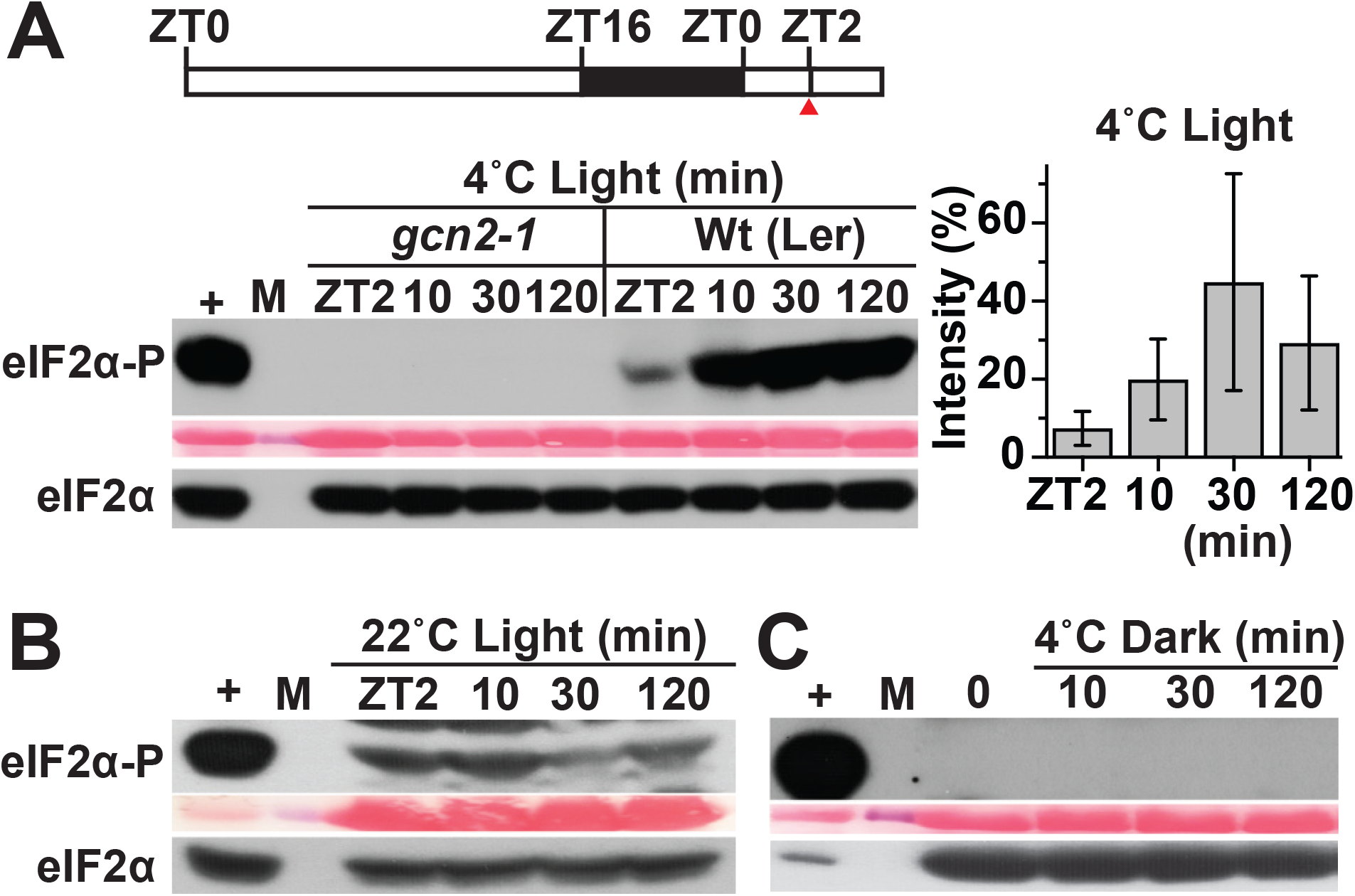
GCN2 kinase activation by cold is light dependent. **(A)** Top - Schematic of the light regimen. Seedlings were grown in a 16 hr light 8 hr dark cycle and shifted to 4°C starting at zeitgeber time (ZT)2. The red arrow at ZT2 indicates the beginning of cold treatment and the start of sampling. Bottom - Immunoblot showing the time course of eIF2*α* phosphorylation in 14-days-old wild-type Landsberg (Wt(Ler)) and *gcn2-1* mutant (*gcn2-1*) seedlings subjected to cold stress as described in panel A. Upper panel: Probed with phospho-specific antibody against eIF2α-P (38kDa). Middle panel: Rubisco large subunit (~ 55kDa) as a loading control after Ponceau S staining of the blot. Lower panel: Probed with antibody against eIF2α (38kDa). (+), arbitrary amount of total protein extract from glyphosate treated Wt seedlings indicating unphosphorylated (eIF2α) or phosphorylated (eIF2α-P) protein; (10, 30, 120) sampling time in minutes; (M) Molecular weight marker. Also shown on the right is the variation in eIF2α-P levels (percent intensity) across the tested time periods in Wt seedlings. Error bars represent Std. deviation from five biological replicates. **(B)** Time course of eIF2α phosphorylation as in panel (A) but with Wt seedlings maintained at 22°C as a control. A partially cropped top band in the eIF2α-P blot indicates non-specific binding of the eIF2α-P antibody. **(C)** eIF2α phosphorylation in Wt seedlings under 4°C in the dark. Seedlings were grown in a 16 hr light 8 hr dark cycle, dark-acclimated for 24 hr and shifted to 4°C in the dark. Time = 0 indicates the beginning of the cold treatment and the start of sampling in dark.

### Salt stress activates GCN2 in a light dependent manner

eIF2α has been shown to get phosphorylated in response to salt stress in mammals (Lu et al., 2001) and yeast (Goossens et al., 2001). To determine this response in plants, Arabidopsis seedlings grown in long-day were shifted to 150mM sodium chloride or an osmotically matched control (300mM mannitol) (Fig. 2A). Similar to other eukaryotes, salt treatment triggered eIF2α phosphorylation within 2 hours only in the wild type but not in the *gcn2-1* mutant seedlings (Fig. 2A). In addition, mock transfer (to 0.1% sucrose) and transfer onto mannitol did not activate GCN2. Similar to cold stress, salt stress too has been linked to adverse effects on chloroplasts in terms of photosynthesis and ROS accumulation (Parida and Das, 2005;Suo et al., 2017;Robles and Quesada, 2019). To test the role of light under salt triggered GCN2 activation, Arabidopsis seedlings were dark adapted for 24 h and shifted to salt or mock (0.1% sucrose) media. Salt treatment in the dark failed to activate GCN2 in wild-type seedlings, similar to the transfer control (Fig. 2B). Taken together, both cold and salt stress require light to activate GCN2.

**Figure 2.**
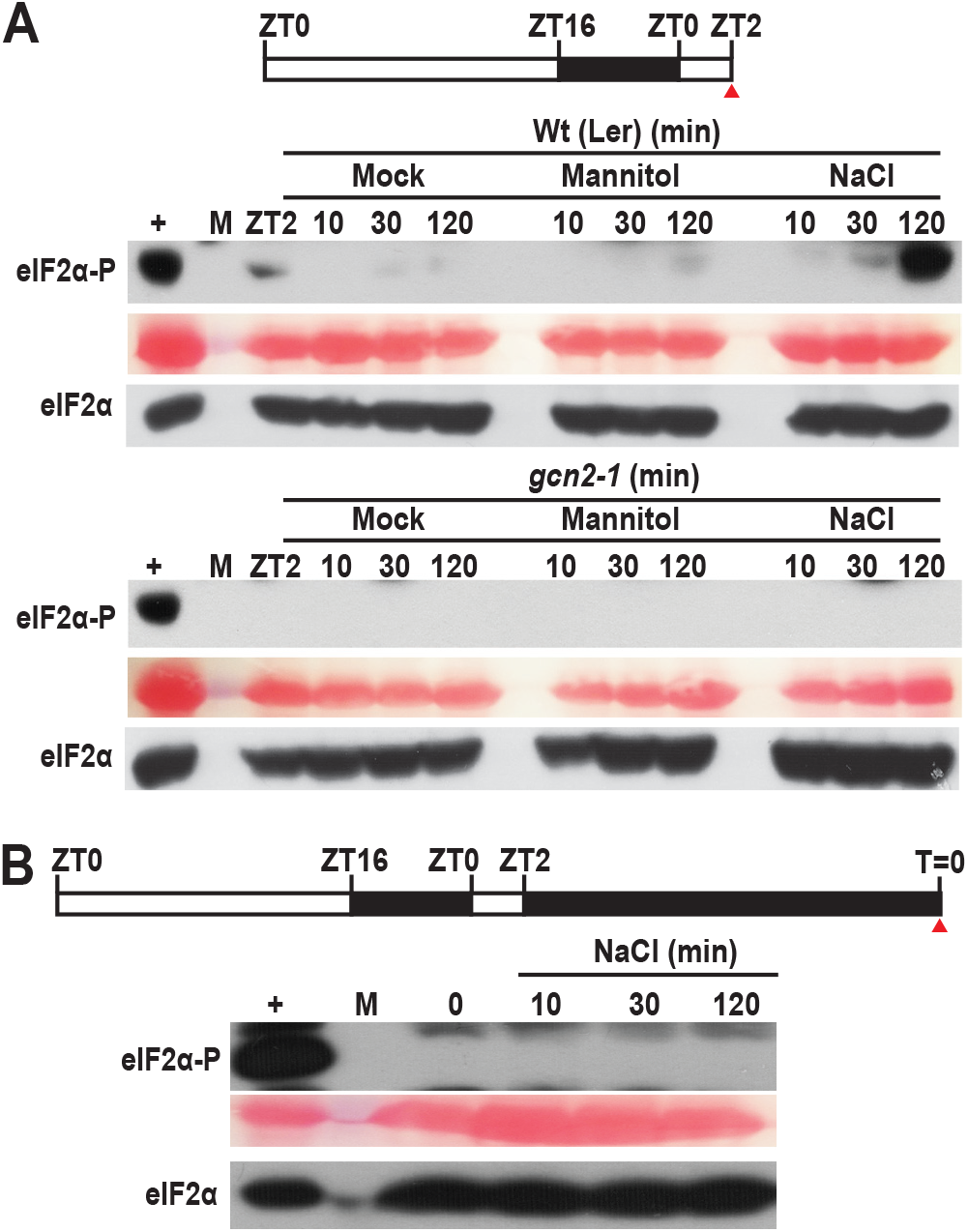
Salt stress activates GCN2 kinase in light. **(A)** Top - Schematic of growth regimen for seedlings under 16hr light and 8hr dark cycle. The red arrow at ZT2=0 indicates the beginning of stress treatment and the start of sampling. Bottom - eIF2 phosphorylation in 10-days-old wild-type Landsberg (Wt (Ler)) and *gcn2-1* mutant (*gcn2-1*) seedlings shifted to media supplemented with 0.1% sucrose (Mock), 150mM Mannitol (Mannitol) or, 150mM NaCl (NaCl). For details see legend to Fig. 1A. **(B)** Top - Schematics of 24 hr dark acclimation starting at ZT2. Bottom - eIF2 phosphorylation in Wt seedlings after shifting to 150mM NaCl in the dark. For details see legend to Fig. 1C.

### Antioxidants and photosynthetic inhibitors alleviate GCN2 activity

In the light, low temperature and salt both affect the photosystem II (PS II), resulting in an increase in the PS II excitation pressure, which generates damaging reactive oxygen species, including hydrogen peroxide (Gray et al., 1996;Huner et al., 1998;Fowler and Thomashow, 2002;Murata et al., 2007). To test whether ROS may contribute to GCN2 activation under cold and salt stress, seedlings were grown in the light on media containing ascorbate and reduced glutathione before challenge with cold or salt stress. These antioxidants delayed the GCN2 activation, albeit weakly in the salt (Fig. 3A), possibly because antioxidants may be barely rate-limiting under these conditions. When eIF2-P was triggered with cold treatment, the presence of ascorbate and glutathione in the medium had only a minor effect (not shown) and there was no detectable boost in H_2_O_2_ levels (Supplemental Figure 1). To address the role of photosynthetic electron transport for GCN2 activity, herbicides that manipulate the plastoquinone (PQ)/ plastoquinol (PQH_2_) pool, 3-(3,4-dichlorophenyl)-1,1-dimethyl urea (DCMU) and 2,5-dibromo-3-methyl-6-isopropyl-p-benzoquinone (DBMIB) were applied shortly prior to the cold and salt treatments. DCMU keeps the plastoquinone pool more oxidized (PQ) and DBMIB more reduced (PQH_2_) (Mateo et al., 2004;Kruk and Karpinski, 2006). Both these herbicides suppressed cold and salt stress triggered GCN2 activation (Fig. 3C, D). These results along with the light dependence of cold and salt stress on GCN2 activation support the notion that chloroplast generated ROS or redox signals may contribute to the activation signal for GCN2, leading to eIF2α phosphorylation.

**Figure 3.**
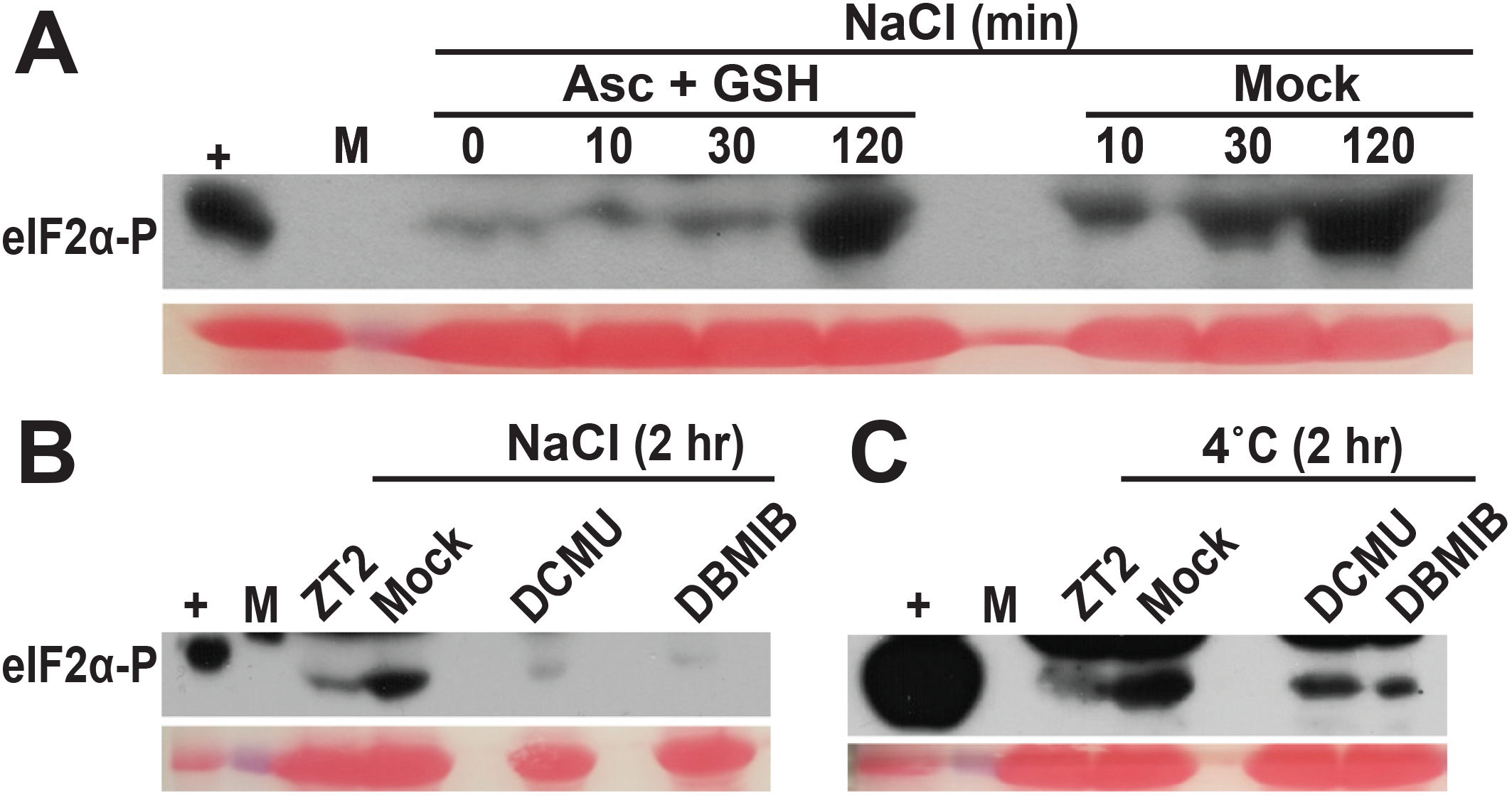
Antioxidant and photosynthetic inhibitors mitigate GCN2 kinase activation under cold and salt stress. **(A)** Time course of eIF2*α* phosphorylation in wild-type Landsberg (Wt) seedlings grown on media supplemented with 0.5mM ascorbate and reduced glutathione for 10-days and shifted to with 150mM NaCl with either antioxidants (Asc + GSH) or mock control. Transfer and sampling of seedlings was performed as described in Fig. 2A. **(B, C)** eIF2α phosphorylation in Wt seedlings treated with either DMSO control (Mock), or 8μM of 3-(3,4-dichlorophenyl)-1,1-dimethylurea (DCMU), or 16μM of 2,5-Dibromo-6-isopropyl-3-methyl-1,4-benzoquinone (DBMIB) thirty minutes prior to **(B)** NaCl or **(C)** 4°C treatment. For details see legend to Fig. 1, 2.

### *gcn2* mutant sensitivity towards cold and salt stress

To determine the role of GCN2 specifically under cold and salt stress conditions at the whole plant level, an established *GCN2* mutant allele (*gcn2-1*) (Lageix et al., 2008;Zhang et al., 2008) in the Landsberg ecotype and a recently characterized homozygous T-DNA insertion allele of *GCN2* in the Columbia ecotype (*gcn2-2)* (Lokdarshi et al., 2019) were tested for phenotypic abnormalities. Under normal growth conditions, *gcn2-1* mutants were indistinguishable from wild type in terms of both shoot and primary root growth (Fig. 4 A, B). However, after challenge with cold stress, *gcn2-1* mutants root lengths were retarded compared to wild type (Fig. 4A, B) as were *gcn2-2* mutants (Supplemental Fig. 2A, B). Of note, the defect in overall growth in the *gcn2* mutants could not be attributed to any defects in the photosynthetic quantum efficiency (Supplemental Figure 3A and B).

**Figure 4.**
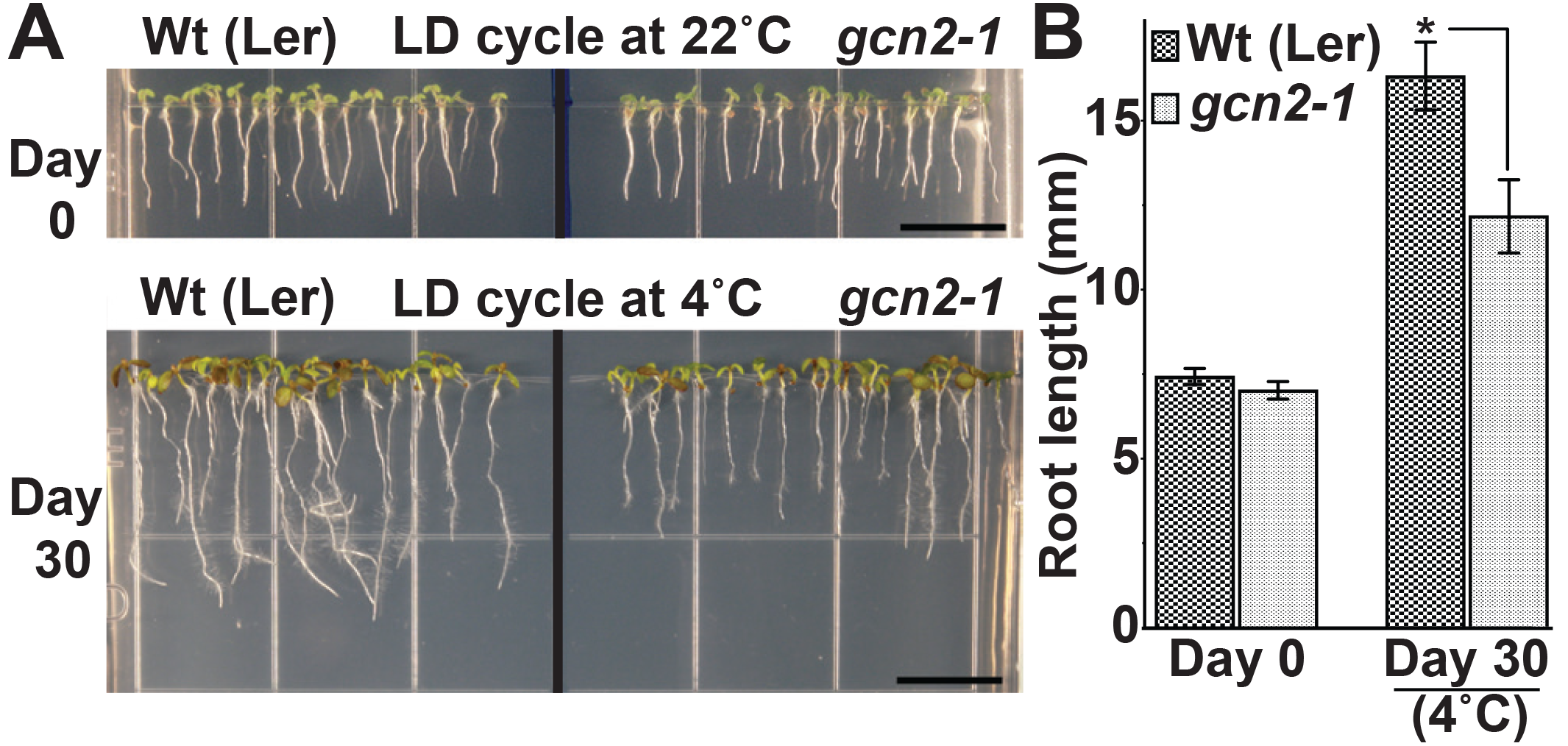
Loss of *GCN2* renders increased sensitivity towards cold stress. **(A)** Top - Representative images of 3-days-old wild-type Landsberg (Wt (Ler)) and *gcn2-1* mutant (*gcn2-1*) seedlings grown under a 16 hr light and 8 hr dark cycle (long day, LD) at 22°C. Seedlings were grown on media with 0.1% sucrose for 3-days and transferred to no sucrose (Day 0). Bottom - Same seedlings after 30 days of LD cycle at 4°C. Scale bars are 10mm. **(B)** Primary root length of Wt and *gcn2-*1 mutant seedlings from panel A. Error bars indicate standard error of the mean from four biological replicates with n>80 per experiment. (Welch’s t-test *P-value <0.05)

Similar to the root growth retardation in the cold, exposure of seedlings to 150 mM NaCl salt also retarded primary root growth in the *gcn2* mutants (Fig. 5A, B; Supplemental Fig. 4A, B). Additionally, *gcn2* mutants showed chlorosis and root growth arrest by day 6 and day 9 (Fig. 5A; Supplemental Fig. 4A: denoted by asterisks). These effects were specific to salt and not seen in the osmotic control (mannitol) and transfer control (0.1% sucrose) treatments. The growth defect of the *gcn2* mutant on salt was evident by Day 6 and resulted in a significant loss of fresh weight and percent survival by Day 9 (Fig. 6A, B). As previously seen for cold stress, the quantum efficiency of PS II declined similarly for *gcn2* and wild type under salt stress (Supplemental Figure 6). We conclude that the *GCN2* gene promotes adaptation of seedlings to cold and salt stress.

**Figure 5.**
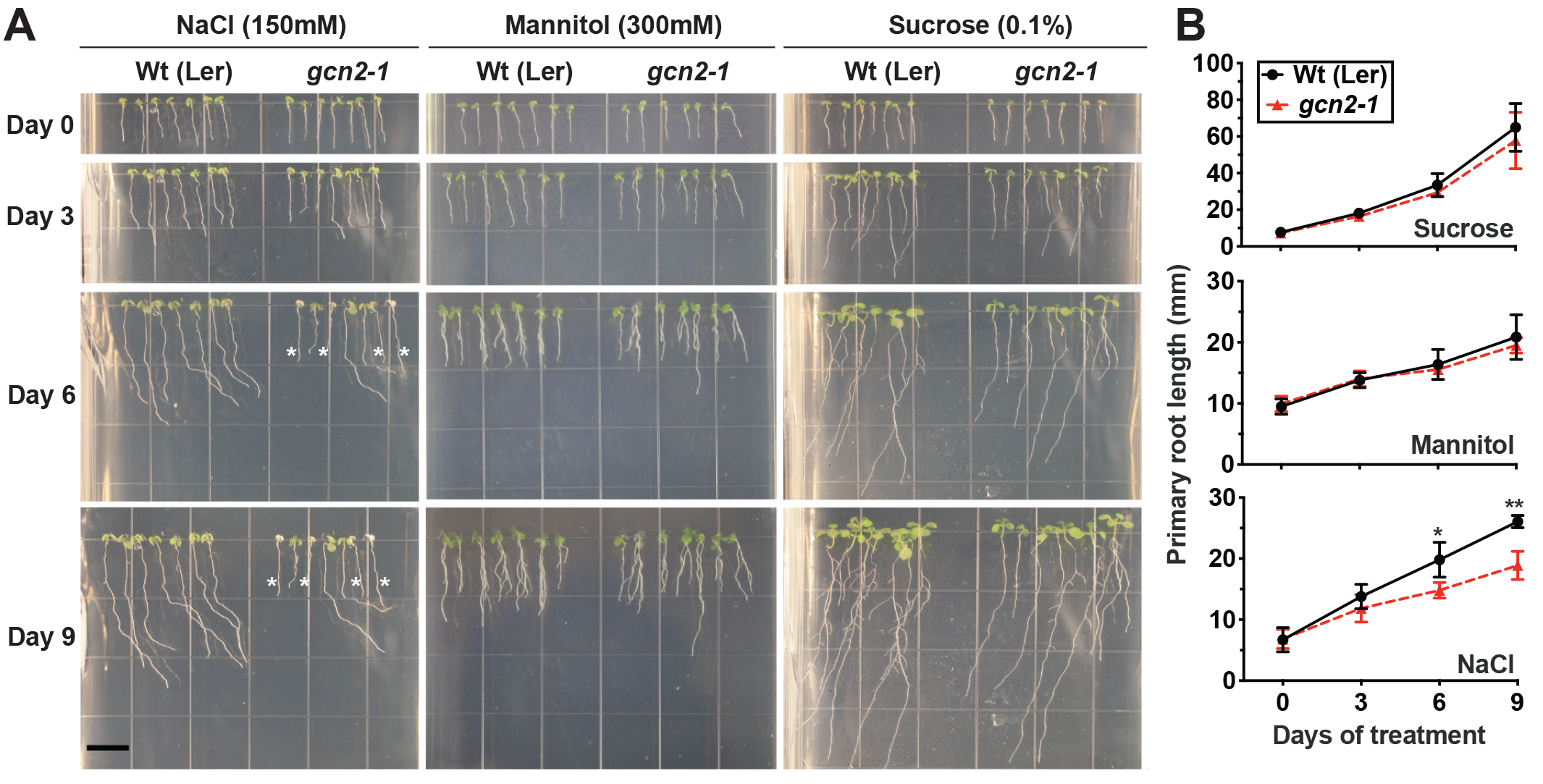
*gcn2* mutants are more sensitive to salt stress. **(A)** Representative images of wild-type Landsberg (Wt (Ler)) and *gcn2-1* mutant (*gcn2-1*) seedlings grown under 16 hr light and 8 hr dark period on plant media supplemented with 150mM NaCl (salt treatment), 300mM Mannitol (osmotic control), or 0.1% sucrose (transfer control). On the day of transfer (Day 0) seedlings were 3 days old on 0.1% sucrose. Scale bar is 10mm. **(B)** Primary root length of Wt and *gcn2-1* mutants from panel (A). Error bars indicate standard error of the mean of four biological replicates with n>36 per experiment (Welch’s t-test *P-value <0.05; ** P-value <0.005).

**Figure 6.**
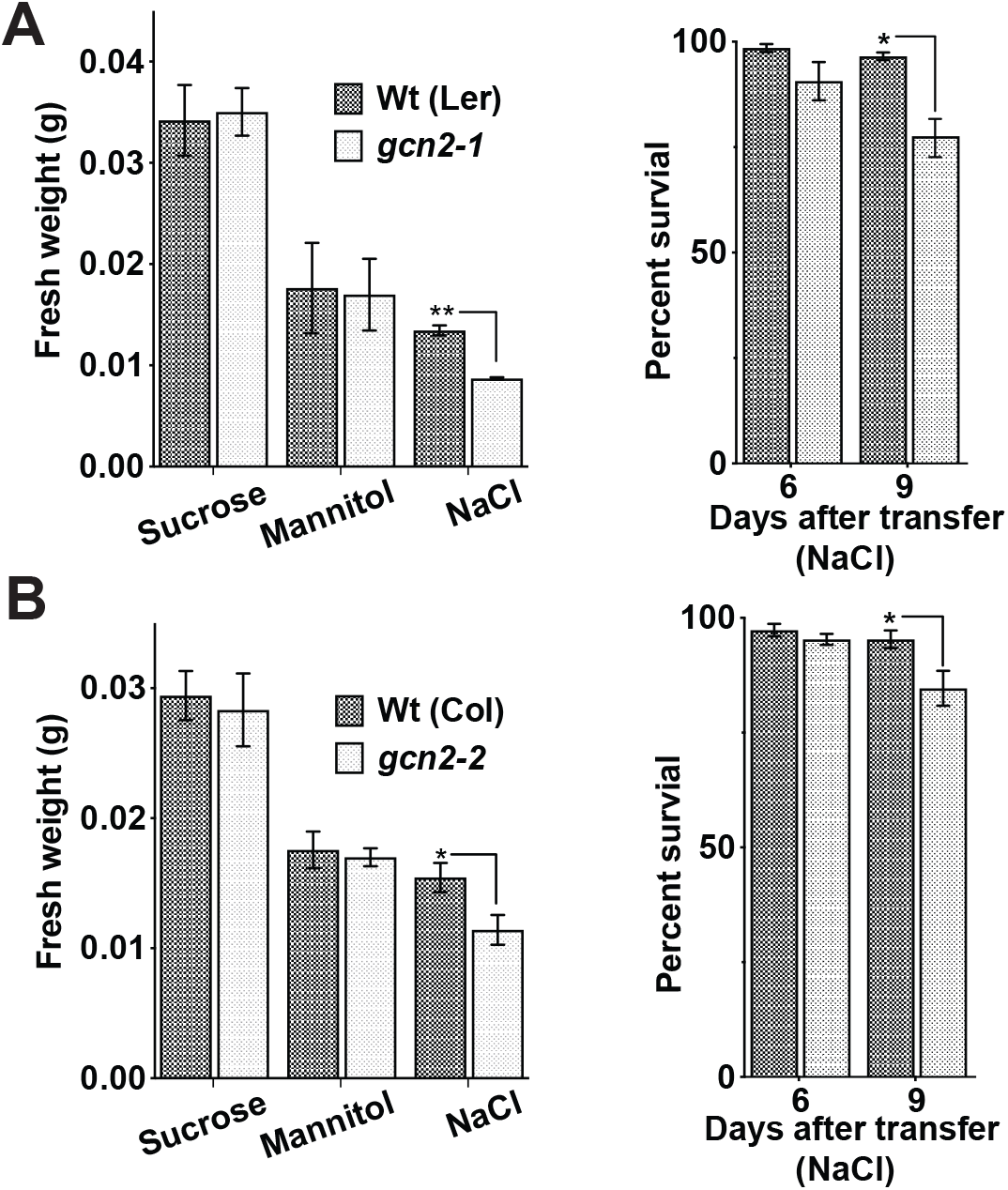
*gcn2* mutants accumulate less fresh weight and exhibit low survival under salt stress. **(A)** Left panel - Fresh weight (grams) of wild-type Landsberg (Wt(Ler)) and *gcn2-1* mutant (*gcn2-* seedlings after 9-days of growth on 0.1% sucrose or, 300mM Mannitol or, 150mM NaCl. Right panel - Percent survival of Wt and *gcn2-1* mutant seedlings at day 6 and day 9 on 150mM NaCl. Analysis performed on seedlings from Fig. 5 **(B)** Fresh weight and percent survival of wild-type Columbia (Wt (Col)) and *gcn2-2* mutant (*gcn2-* seedlings as described in panel A. Analysis performed on seedlings from Supplemental Fig. 4 Error bars indicate standard error of the mean of four biological replicates with n>36 per experiment (Welch’s t-test *P-value <0.05; **P-value <0.005; *** P-value <0.0005).

### mRNA-ribosome loading under cold and salt stress

GCN2 activity has been implicated in the down-regulation of mRNA translation under a variety of stress conditions (Lageix et al., 2008;Zhang et al., 2008;Liu et al., 2015a;Wang et al., 2017). To test the role of GCN2 in global mRNA translation under cold and salt stress, *gcn2* mutant and wild-type seedlings were challenged with the respective stresses under light. Polysome profiles from sucrose density gradients revealed overall similar profiles and polysome-to-monosome ratios for wild-type and *gcn2* under both normal growth conditions (Fig. 7A) and after cold stress (Fig. 7B, C). Likewise, in response to salt stress, both wild-type and *gcn2* mutant displayed similar polysome profiles (Fig. 8). The trend towards slightly elevated ribosome loading in *gcn2-1*, while not uncommon, was not statistically significant. The lack of a clear effect on global polyribosome loading stands in contrast to data after herbicide treatment where ribosome loading declines in a GCN2-dependent manner (Lageix et al., 2008;Lokdarshi et al., 2019).

**Figure 7.**
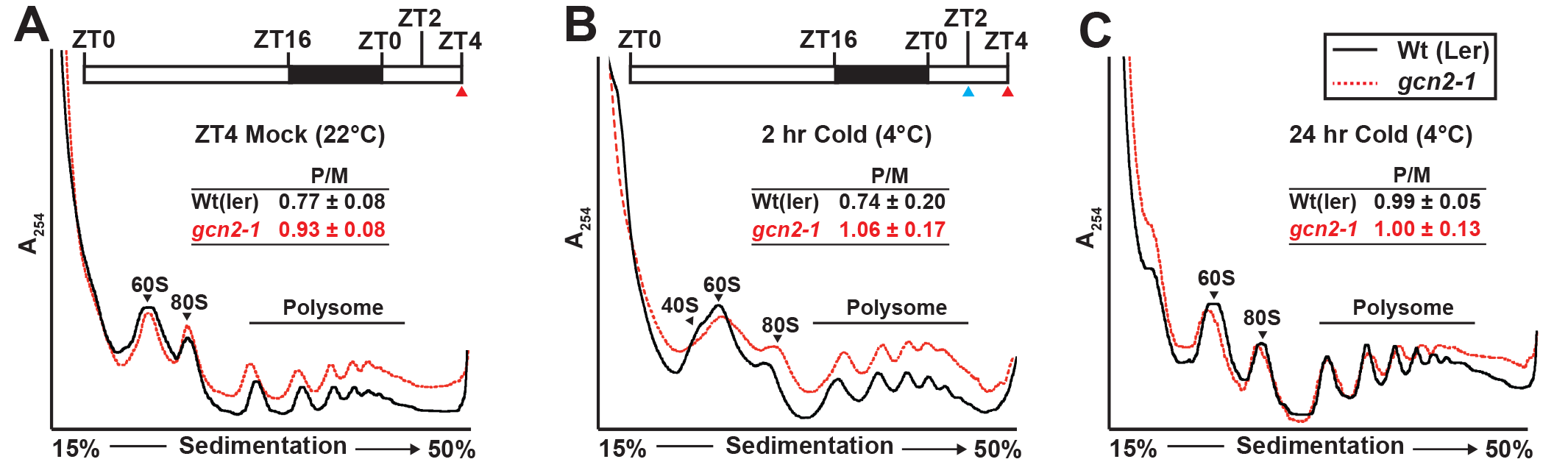
Ribosome-RNA profile of wild-type and *gcn2-1* under standard growth conditions and cold stress. Top - **(A, B)** Schematic of light regimen showing seedling growth in long day period (16 hr light and 8 hr dark) indicating the beginning of cold (4°C) treatment starting at ZT2 (blue arrow) and the sampling time at ZT4 (red arrow). Bottom - UV Absorbance profile at 254nm of 14-days-old wild-type Landsberg (Wt(ler)) and *gcn2-1* mutant (*gcn2-1*) seedlings at (A) 22°C at ZT4 (Mock) or subjected to cold at 4°C **(B)** for 2 hr, or **(C)** for 24 hr under a long day period. The positions of the 40S, 60S, 80S and the polysomes are indicated on the profiles. The ratio of polysomes (P) to monosomes (M) is indicated with standard error from 3 replicates.

**Figure 8.**
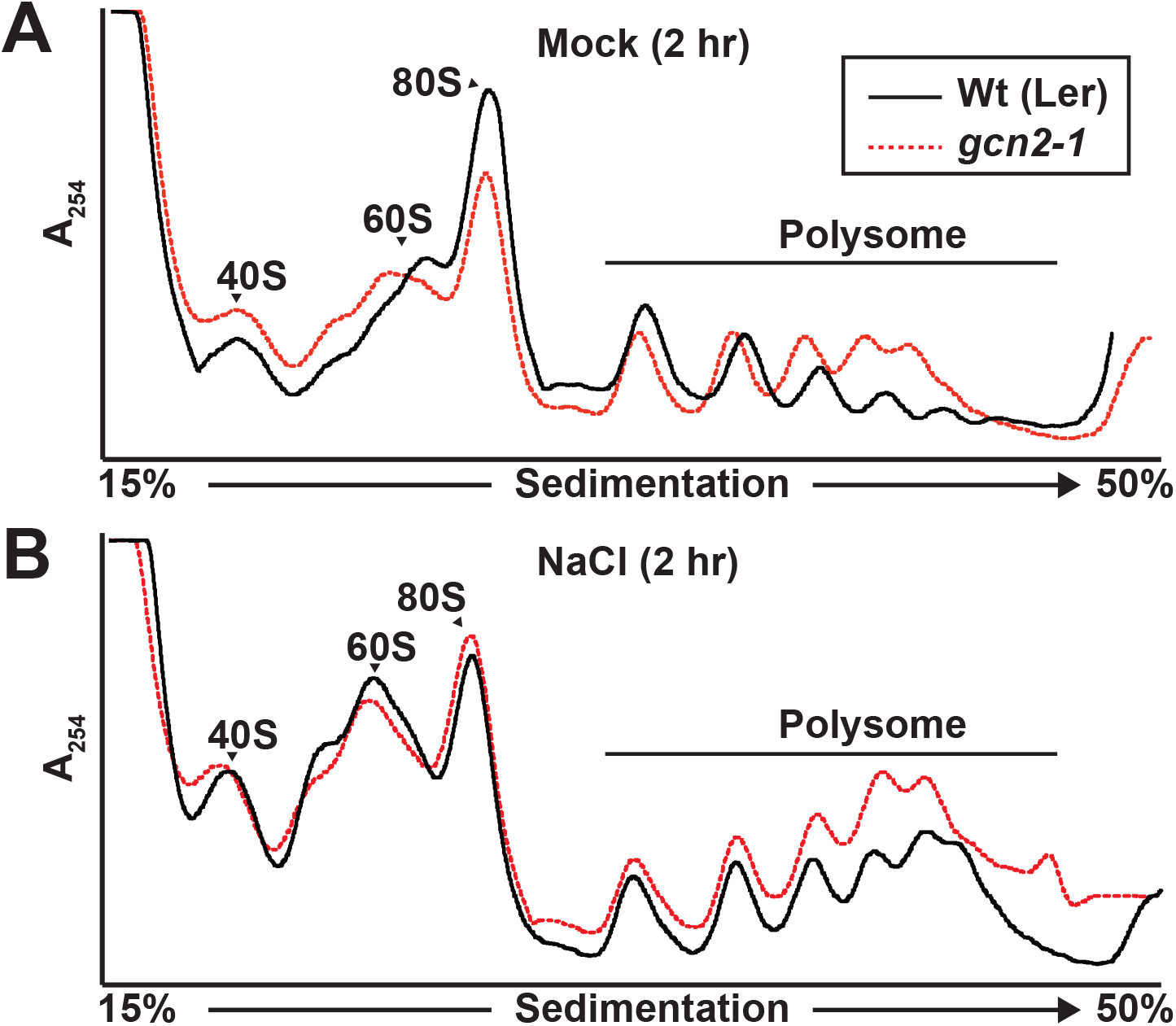
Ribosome-RNA profile of wild-type and *gcn2* mutant under salt stress. Representative UV absorbance (A254) profile of 10-days-old Wild-type Landsberg (Wt(Ler)) and *gcn2-1* mutant (*gcn2-1*) seedlings after 2 hr of treatment with **(A)** 0.1% sucrose (Mock) or **(B)** NaCl. Seedling transfer was performed as described in Figure 2. Positions of the 40S, 60S, 80S and the polysome are indicated on the profile.

## DISCUSSION

The GCN2-eIF2α module is an integral component of a pan-eukaryotic stress response program. In yeast and mammals, GCN2 is activated by binding to uncharged tRNAs via its C-terminal HisRS domain. In plants, GCN2 kinase is activated under a wide range of abiotic stresses (e.g., UV light, cold, wounding), synthetic agents (e.g., herbicides, purine starvation), hormones (e.g., methyl jasmonate, salicylic acid, abscisic acid) and live bacterial pathogen (e.g., *Pseudomonas syringae*). More recently, Arabidopsis GCN2 was found to be activated in response to H_2_O_2_ directly, as well as excess light stress and methyl viologen, treatments that produce ROS (Lokdarshi et al., 2019). In the present study, we show that both cold and salt challenge not only activate eIF2α-P but require light to do so, similar to our recent findings of GCN2 activation in response to herbicide. Taken together, our study suggests that the highly conserved GCN2-eIF2α module is activated in a common manner by different stresses, possibly by ROS, given that H_2_O_2_ is the only known signal to activate GCN2 in darkness (Lokdarshi et al., 2019). The precise biochemical mechanism remains to be determined.

Biochemically, the only known ligand to activate plant GCN2 *in vitro* are uncharged tRNAs, which presumably accumulate in the cell during amino acid starvation. Whether uncharged tRNAs are necessary and sufficient to activate GCN2 *in planta* under all stress conditions remains unclear. It is plausible that tRNA is bound to GCN2 as a coactivator but that additional signals are needed to boost kinase activity to physiologically relevant levels. Of note, recently Inglis and coworkers reported that mammalian GCN2 can be activated in a tRNA-independent mechanism by the ribosomal P-stalk protein complex (Sattlegger and Hinnebusch, 2000;Inglis et al., 2019). The mechanism of how GCN2 is activated *in planta* by tRNAs and ROS may also depend on the GCN2 interacting proteins GCN1 and GCN20 (Wang et al., 2017; Izquierdo et al., 2018), similar to yeast and mammals; however plastidic ROS as a GCN2 activation signal is unique to plants.

It remains unclear whether and how the GCN2 mediated phosphorylation of eIF2α under various conditions drives global translational repression as seen at the level of polyribosome loading, and how this response supports plant growth and development. The clearest causal chain of events is observed with herbicides that inhibit amino acid synthesis, where activation of GCN2 kinase by herbicide in the presence of light-conditioned ROS causes eIF2α phosphorylation, followed by global translational repression, which is disrupted in the *gcn2* mutant (Lageix et al., 2008;Lokdarshi et al., 2019). Moreover, the *gcn2* mutant is hypersensitive to herbicide (Zhang et al., 2008;Izquierdo et al., 2018), all in keeping with a simple, linear signaling pathway. However, it is much less clear how other GCN2-targeted abiotic stimuli affect translation, notwithstanding that it has been confirmed multiple times that eIF2α phosphorylation is always mediated by GCN2. Here we showed that upon cold treatment, eIF2α became phosphorylated by GCN2, but with no detectable translational repression by either cold or GCN2 kinase, although *gcn2* mutants were cold sensitive. We observed the same result for salt stress. Of note, salt stress at slightly higher intensity in rice (Ueda et al., 2012), but not cold stress in Arabidopsis (Juntawong et al., 2013) cause a drop in global ribosome loading. As for ROS, which we consider the most immediate activator of the GCN2 kinase, this stress represses translation as well as plant growth, but neither is detectably GCN2-dependent (Lokdarshi et al., 2019). The same pattern was seen in response to DTT and antimycinA (Izquierdo et al., 2018). Under high light, which is likely another relevant trigger of GCN2 in the natural environment, again, there is no GCN2-dependent translational repression, although *gcn2* mutants are sensitive to high light (Lokdarshi et al., 2019). For comparison, heat and hypoxia both rapidly repress global translation (Branco-Price et al., 2008;Matsuura et al., 2010;Yanguez et al., 2013), but without any apparent phosphorylation of eIF2α. Taken together, these observations clearly suggest that, despite the seemingly simple sequence of events in response to certain inhibitors of amino acid synthesis, not every instance of eIF2α phosphorylation causes global translational repression, and only some but not all instances of global translational repression are conditioned on eIF2α phosphorylation. These observations indicate that there must be additional translational control pathways that cooperate with GCN2-mediated eIF2α phosphorylation to organize the translatome under abiotic stress. Candidates are GCN1 and GCN20-mediated (Wang et al., 2017; Izquierdo et al., 2018), and autophagy-mediated processes (Zhao et al., 2018; Yoon and Chung, 2019) and processes involving SnRK-TOR signaling (Margalha et al., 2019) and stress granules (Chantarachot and Bailey-Serres, 2018). This conclusion is also in keeping with the emerging role of GCN2 in responses to plant pathogens. Under certain conditions, pathogens or effectors of immunity activate GCN2 or eIF2α phosphorylation (Pajerowska-Mukhtar et al., 2012;Liu et al., 2019) while in other conditions they do not (Zhang et al., 2008;Meteignier et al., 2017;Izquierdo et al., 2018). Certain pathogens do trigger translational reorganizations (Moeller et al., 2012;Xu et al., 2017) and GCN2 is involved in responses to bacterial pathogens (Liu et al., 2015b;Liu et al., 2019;Lokdarshi et al., 2019) although the precise role of GCN2 kinase signaling in defense related translational control remains to be defined.

Overall, the findings presented in this study add to a unified model of the regulation of the cytosolic translation apparatus via the highly conserved GCN2-eIF2α module under a variety of abiotic stresses, that may also extend to biotic stresses in plants. In summary, we show that activation of GCN2 by cold and salt stress is dependent on the redox state of the chloroplast, and loss of *GCN2* results in the increased sensitivity towards common abiotic stress inputs, cold and salt. In the future, determining what biochemical and molecular events lead to GCN2 activation under these natural stress inputs will shed light on the integrated stress response pathway in plants. Additionally, the regulation of global translation versus specific mRNAs that fall under stress type regulation is also a subject of further investigation.

## ACKNOWLEDGMENTS

We thank Dr. Karen Browning for the antibody against eIF2α and Ricardo Urquidi-Camacho for assistance with data analysis and discussion.

## AUTHOR CONTRIBUTIONS

AL, PM, MF, ZE and CE performed the experiments

AL, AVA – Analyzed the results and wrote the manuscript

## SUPPLEMENTARY MATERIAL

**Supplemental Figure 1. Cold treatment barely modulates hydrogen peroxide levels.**

**Supplemental Figure 2. Loss of *GCN2* renders increased sensitivity towards cold stress in the Columbia ecotype.**

**Supplemental Figure 3. Effect of cold stress on photosynthetic efficiency of wild-type and *gcn2* mutants.**

**Supplemental Figure 4. Loss of *GCN2* renders increased sensitivity towards salt stress in the Columbia ecotype**.

**Supplemental Figure 5. Effect of salt stress on photosynthetic efficiency of wild-type and *gcn2.***

## REFERENCES

Adam, S., and Murthy, S.D.S. (2014). “Effect of Cold Stress on Photosynthesis of Plants and Possible Protection Mechanisms,” in Approaches to Plant Stress and their Management, eds. R.K. Gaur & P. Sharma. (New Delhi: Springer India), 219–226.

Anda, S., Zach, R., and Grallert, B. (2017). Activation of Gcn2 in response to different stresses. PLoS One 12, e0182143.

Benina, M., Ribeiro D.M, Gechev T.S, Mueller-Roeber, B. and Schippers J.H (2015). A cell type-specific view on the translation of mRNAs from ROS-responsive genes upon paraquat treatment of Arabidopsis thaliana leaves. Plant Cell Env 38, 349–363.

Branco-Price, C., Kaiser K.A, Jang C.J, Larive C.K, and Bailey-Serres, J. (2008). Selective mRNA translation coordinates energetic and metabolic adjustments to cellular oxygen deprivation and reoxygenation in Arabidopsis thaliana. Plant J 56, 743–755.

Chantarachot, T., and Bailey-Serres, J. (2018). Polysomes, Stress Granules, and Processing Bodies: A Dynamic Triumvirate Controlling Cytoplasmic mRNA Fate and Function. Plant Physiol 176 254–269.

Crosatti, C., Rizza, F., Badeck F.W, Mazzucotelli, E., and Cattivelli, L. (2013). Harden the chloroplast to protect the plant. Physiologia Plantarum 147 55–63.

Dennis M.D, Person M.D, and Browning K.S (2009). Phosphorylation of plant translation initiation factors by CK2 enhances the in vitro interaction of multifactor complex components. J Biol Chem 284 20615–20628.

Dever T.E, Feng, L., Wek R.C, Cigan A.M, Donahue T.F, and Hinnebusch A.G (1992). Phosphorylation of initiation factor 2 alpha by protein kinase GCN2 mediates gene-specific translational control of GCN4 in yeast. Cell 68 585–596.

Dong, J., Qiu, H., Garcia-Barrio, M., Anderson, J., and Hinnebusch A.G (2000). Uncharged tRNA activates GCN2 by displacing the protein kinase moiety from a bipartite tRNA-binding domain. Mol Cell 6 269–279.

Donnelly, N., Gorman A.M, Gupta, S., and Samali, A. (2013). The eIF2alpha kinases: their structures and functions. Cell Mol Life Sci 70 3493–3511.

Enganti, R., Cho S.K, Toperzer J.D, Urquidi-Camacho, R.A., Cakir O.S, Ray A.P, Abraham P.E, Hettich R.L, and von Arnim, A.G. (2018). Phosphorylation of Ribosomal Protein RPS6 Integrates Light Signals and Circadian Clock Signals. Frontiers in Plant Science 8, 2210.

Faus, I., Zabalza, A., Santiago, J., Nebauer S.G, Royuela, M., Serrano, R., and Gadea, J. (2015). Protein kinase GCN2 mediates responses to glyphosate in Arabidopsis. BMC Plant Biol 15, 14.

Fowler, S., and Thomashow M.F (2002). Arabidopsis transcriptome profiling indicates that multiple regulatory pathways are activated during cold acclimation in addition to the CBF cold response pathway. Plant Cell 14 1675–1690.

Goossens, A., Dever T.E, Pascual-Ahuir, A., and Serrano, R. (2001). The protein kinase Gcn2p mediates sodium toxicity in yeast. J Biol Chem 276 30753–30760.

Gray G.R, Savitch L.V, Ivanov A.G, and Huner, N. (1996). Photosystem II Excitation Pressure and Development of Resistance to Photoinhibition (II. Adjustment of Photosynthetic Capacity in Winter Wheat and Winter Rye). Plant Physiol 110 61–71.

Harding H.P, Novoa, I., Zhang, Y., Zeng, H., Wek, R., Schapira, M., and Ron, D. (2000). Regulated translation initiation controls stress-induced gene expression in mammalian cells. Mol Cell 6 1099–1108.

Hinnebusch A.G, Ivanov I.P, and Sonenberg, N. (2016). Translational control by 5’-untranslated regions of eukaryotic mRNAs. Science 352 1413–1416.

Huner, N.P.A., Oquist, G., and Sarhan, F. (1998). Energy balance and acclimation to light and cold. Trends in Plant Science 3 224–230.

Inglis A.J, Masson G.R, Shao, S., Perisic, O., Mclaughlin S.H, Hegde R.S, and Williams R.L (2019). Activation of GCN2 by the ribosomal P-stalk. Proc Natl Acad Sci U S A 116 4946–4954.

Izquierdo, Y., Kulasekaran, S., Benito, P., Lopez, B., Marcos, R., Cascon, T., Hamberg, M., and Castresana, C. (2018). Arabidopsis nonresponding to oxylipins locus NOXY7 encodes a yeast GCN1 homolog that mediates noncanonical translation regulation and stress adaptation. Plant Cell and Environment 41 1438–1452.

Juntawong, P., and Bailey-Serres, J. (2012). Dynamic light regulation of translation status in Arabidopsis thaliana. Frontiers in Plant Science 3, 66.

Juntawong, P., Sorenson, R., and Bailey-Serres, J. (2013). Cold shock protein 1 chaperones mRNAs during translation in Arabidopsis thaliana. Plant J 74 1016–1028.

Kashiwagi, K., Yokoyama, T., Nishimoto, M., Takahashi, M., Sakamoto, A., Yonemochi, M., Shirouzu, M., and Ito, T. (2018). Structural Basis for eIF2B Inhibition in Integrated Stress Response. Science 364 495–499.

Khandal, D., Samol, I., Buhr, F., Pollmann, S., Schmidt, H., Clemens, S., Reinbothe, S., and Reinbothe, C. (2009). Singlet oxygen-dependent translational control in the tigrina-d.12 mutant of barley. Proceedings of the National Academy of Sciences of the United States of America 106, 13112–13117.

Kruk, J., and Karpinski, S. (2006). An HPLC-based method of estimation of the total redox state of plastoquinone in chloroplasts, the size of the photochemically active plastoquinone-pool and its redox state in thylakoids of Arabidopsis. Biochim Biophys Acta 1757 1669–1675.

Lageix, S., Lanet, E., Pouch-Pelissier, M.N., Espagnol M.C, Robaglia, C., Deragon J.M, and Pelissier, T. (2008). Arabidopsis eIF2alpha kinase GCN2 is essential for growth in stress conditions and is activated by wounding. BMC Plant Biol 8, 134.

Li M.W, Auyeung W.K, and Lam H.M (2013). The GCN2 homologue in Arabidopsis thaliana interacts with uncharged tRNA and uses Arabidopsis eIF2alpha molecules as direct substrates. Plant Biol (Stuttg) 15 13–18.

Liu, B., and Qian S.B (2014). Translational reprogramming in cellular stress response. Wiley Interdiscip Rev RNA 5 301–315.

Liu M.J, Wu S.H, Chen H.M, and Wu S.H (2012). Widespread translational control contributes to the regulation of Arabidopsis photomorphogenesis. Molecular Systems Biology 8, 566.

Liu, X., Afrin, T., and Pajerowska-Mukhtar, K.M. (2019). Arabidopsis GCN2 kinase contributes to ABA homeostasis and stomatal immunity. Communications Biology 2, 302.

Liu, X., Kørner, C.J., Hajdu, D., Guo, T., Ramonell K.M, Argueso C.T, and Pajerowska-Mukhtar,K.M. (2015a). Arabidopsis Thaliana Atgcn2 Kinase Is Involved In Disease Resistance Against Pathogens With Diverse Life Styles. International Journal of Phytopathology 4, 12.

Liu, X., Merchant, A., Rockett K.S, Mccormack, M., and Pajerowska-Mukhtar, K.M. (2015b). Characterization of Arabidopsis thaliana GCN2 kinase roles in seed germination and plant development. Plant Signal Behav 10, e992264.

Lokdarshi, A., Conner W.C, Mcclintock, C., Li, T., and Roberts D.M (2016). Arabidopsis CML38, a Calcium Sensor That Localizes to Ribonucleoprotein Complexes under Hypoxia Stress. Plant Physiol 170 1046–1059.

Lokdarshi, A., Guan, J., Urquidi Camacho R.A, Cho S.K, Morgan P.W, Leonard, M., Shimono, M., Day, B., and von Arnim, A.G. (2019). Light activates the translational regulatory GCN2 kinase via reactive oxygen species emanating from the chloroplast. bioRxiv, 794362.

Lu, L., Han A.P, and Chen J.J (2001). Translation initiation control by heme-regulated eukaryotic initiation factor 2alpha kinase in erythroid cells under cytoplasmic stresses. Mol Cell Biol 21 7971–7980.

Margalha, L., Confraria, A., and Baena-González, E. (2019). SnRK1 and TOR: modulating growth#8211; defense trade-offs in plant stress responses. Journal of Experimental Botany 70 2261–2274.

Mateo, A., Muhlenbock, P., Rusterucci, C., Chang C.C, Miszalski, Z., Karpinska, B., Parker J.E, Mullineaux P.M, and Karpinski, S. (2004). LESION SIMULATING DISEASE 1 is required for acclimation to conditions that promote excess excitation energy. Plant Physiol 136 2818–2830.

Matsuura, H., Kiyotaka, U., Ishibashi, Y., Kubo, Y., Yamaguchi, M., Hirata, K., Demura, T., and Kato,K. (2010). A short period of mannitol stress but not LiCl stress led to global translational repression in plants. Biosci Biotechnol Biochem 74 2110–2112.

Merchante, C., Stepanova A.N, and Alonso J.M (2017). Translation regulation in plants: an interesting past, an exciting present and a promising future. Plant J 90 628–653.

Meteignier L.V, El Oirdi, M., Cohen, M., Barff, T., Matteau, D., Lucier J.F, Rodrigue, S., Jacques P.E, Yoshioka, K., and Moffett, P. (2017). Translatome analysis of an NB-LRR immune response identifies important contributors to plant immunity in Arabidopsis. J Exp Bot 68, 2333–2344.

Missra, A., Ernest, B., Lohoff, T., Jia, Q., Satterlee, J., Ke, K., and Von Arnim, A.G. (2015). The Circadian Clock Modulates Global Daily Cycles of mRNA Ribosome Loading. Plant Cell 27 2582–2599.

Moeller J.R, Moscou M.J, Bancroft, T., Skadsen R.W, Wise R.P, and Whitham S.A (2012). Differential accumulation of host mRNAs on polyribosomes during obligate pathogen-plant interactions. Mol Biosyst 8 2153–2165.

Murata, N., Takahashi, S., Nishiyama, Y., and Allakhverdiev S.I (2007). Photoinhibition of photosystem II under environmental stress. Biochim Biophys Acta 1767 414–421.

Murchie E.H, and Lawson, T. (2013). Chlorophyll fluorescence analysis: a guide to good practice and understanding some new applications. J Exp Bot 64 3983–3998.

Pajerowska-Mukhtar, K.M., Wang, W., Tada, Y., Oka, N., Tucker C.L, Fonseca J.P and Dong, X. (2012). The HSF-like transcription factor TBF1 is a major molecular switch for plant growth-to-defense transition. Curr Biol 22 103–112.

Parida A.K, and Das A.B (2005). Salt tolerance and salinity effects on plants: a review. Ecotoxicol Environ Saf 60 324–349.

Robles, P., and Quesada, V. (2019). Transcriptional and Post-transcriptional Regulation of Organellar Gene Expression (OGE) and Its Roles in Plant Salt Tolerance. Int J Mol Sci 20, pii: E1056.

Sattlegger, E., and Hinnebusch A.G (2000). Separate domains in GCN1 for binding protein kinase GCN2 and ribosomes are required for GCN2 activation in amino acid-starved cells. The EMBO Journal 19 6622–6633.

Suo, J., Zhao, Q., David, L., Chen, S., and Dai, S. (2017). Salinity Response in Chloroplasts: Insights from Gene Characterization. Int J Mol Sci 18, pii: E1011.

Tang, L., Bhat, S., and Petracek M.E (2003). Light control of nuclear gene mRNA abundance and translation in tobacco. Plant Physiol 133 1979–1990.

Ueda, K., Matsuura, H., Yamaguchi, M., Demura, T., and Kato, K. (2012). Genome-wide analyses of changes in translation state caused by elevated temperature in Oryza sativa. Plant Cell Physiol 53 1481–1491.

Wang, L., Li, H., Zhao, C., Li, S., Kong, L., Wu, W., Kong, W., Liu, Y., Wei, Y., Zhu J.K, and Zhang, H. (2017). The inhibition of protein translation mediated by AtGCN1 is essential for cold tolerance in Arabidopsis thaliana. Plant Cell Environ 40 56–68.

Wek S.A, Zhu, S., and Wek R.C (1995). The histidyl-tRNA synthetase-related sequence in the eIF-2 alpha protein kinase GCN2 interacts with tRNA and is required for activation in response to starvation for different amino acids. Mol Cell Biol 15 4497–4506.

Xu, G., Greene G.H, Yoo, H., Liu, L., Marqués, J., Motley, J., and Dong, X. (2017). Global translational reprogramming is a fundamental layer of immune regulation in plants. Nature 545, 487.

Yanguez, E., Castro-Sanz, A.B., Fernandez-Bautista, N., Oliveros J.C, and Castellano M.M (2013). Analysis of genome-wide changes in the translatome of Arabidopsis seedlings subjected to heat stress. PLoS One 8, e71425.

Yoon S.H, and Chung, T. (2019). Protein and RNA Quality Control by Autophagy in Plant Cells. Molecules and cells 42 285–291.

Zhang, Y., Dickinson J.R, Paul M.J, and Halford N.G (2003). Molecular cloning of an Arabidopsis homologue of GCN2, a protein kinase involved in co-ordinated response to amino acid starvation. Planta 217 668–675.

Zhang, Y., Wang, Y., Kanyuka, K., Parry M.A, Powers S.J, and Halford N.G (2008). GCN2-dependent phosphorylation of eukaryotic translation initiation factor-2alpha in Arabidopsis. J Exp Bot 59 3131–3141.

Zhao, L., Deng, L., Zhang, Q., Jing, X., Ma, M., Yi, B., Wen, J., Ma, C., Tu, J., Fu, T., and Shen, J. (2018). Autophagy contributes to sulfonylurea herbicide tolerance via GCN2-independent regulation of amino acid homeostasis. Autophagy 14 702–714.

