## Supplemental Figure 1 for "Light dependent activation of the GCN2 kinase under cold and salt stress is mediated by the photosynthetic status of the chloroplast"

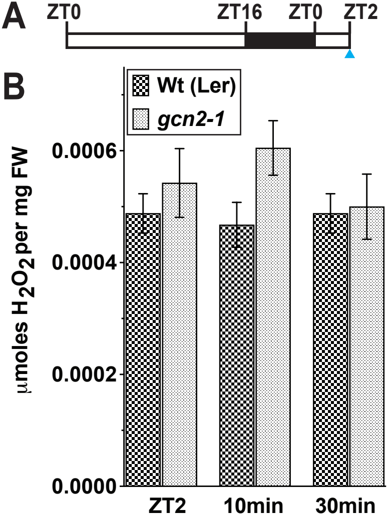


**Supplemental Figure 1. Cold treatment barely modulates hydrogen peroxide levels.**
**(A)** Light regimen showing the growth of seedlings in 16 hr light and 8 hr dark cycle and the start of cold treatment at ZT2 (blue arrow).
**(B)** Relative $H_2O_2$ levels in 14-days-old wild-type Landsberg (Wt (Ler)) and *gcn2-1* mutant *(gcn2-1)* seedlings at ZT2, and after 10 min and 30 min of cold treatment. Error bars represent standard error of the mean of 8 biological replicates. Welch's *t*-test *P*-values were >0.5.
