## Supplemental Figure 2 for "Light dependent activation of the GCN2 kinase under cold and salt stress is mediated by the photosynthetic status of the chloroplast"

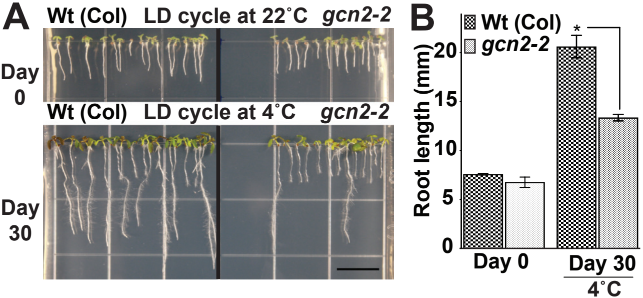


**Supplemental Figure 2. Loss of *GCN2* renders increased sensitivity towards cold stress in the Columbia ecotype.**
**(A)** Top - Representative images of 3-days-old wild-type Columbia (Wt (Col)) and *gcn2-2* mutant (*gcn2-2*) seedlings grown under 16 hr light and 8 hr dark long day (LD) cycle at 22°C. Seedlings were grown on media with 0.1% sucrose for 3-days and transferred to no sucrose (Day 0).
Bottom - Same seedlings after 30 days of LD cycle at 4°C. Scale bars are 10mm.
**(B)** Primary root length of Wt and *gcn2-2* mutant seedlings from panel A. Error bars indicate standard error of the mean from four biological replicates with n>80 per experiment. (Welch's t-test *P-value <0.05)
