## Supplemental Figure 3 for "Light dependent activation of the GCN2 kinase under cold and salt stress is mediated by the photosynthetic status of the chloroplast"

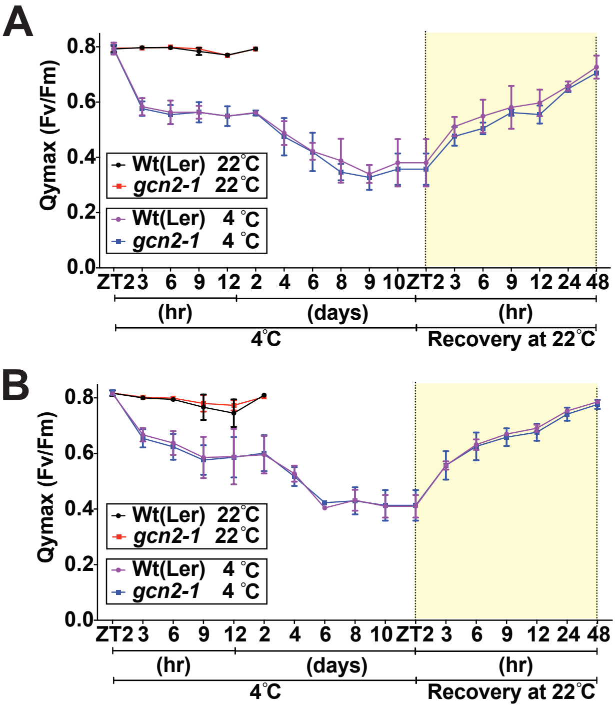


**Supplemental Figure 3 . Effect of cold stress on photosynthetic efficiency of wild-type and *gcn2* mutants.** Time course analysis of photosystem II maximal quantum yield (Qymax (Fv/Fm)) in rosette stage plants of **(A)** wild-type Landsberg (Wt (Ler)) and *gcn2-1* mutant , **(B)** wild-type Columbia (Wt(Col)) and *gcn2-2* mutants under cold stress at 4°C and recovery at 22°C. Error bars represent standard error of the mean from three biological replicates. Welch's t-test *P*-value >0.5.
