## Supplemental Figure 4 for "Light dependent activation of the GCN2 kinase under cold and salt stress is mediated by the photosynthetic status of the chloroplast"

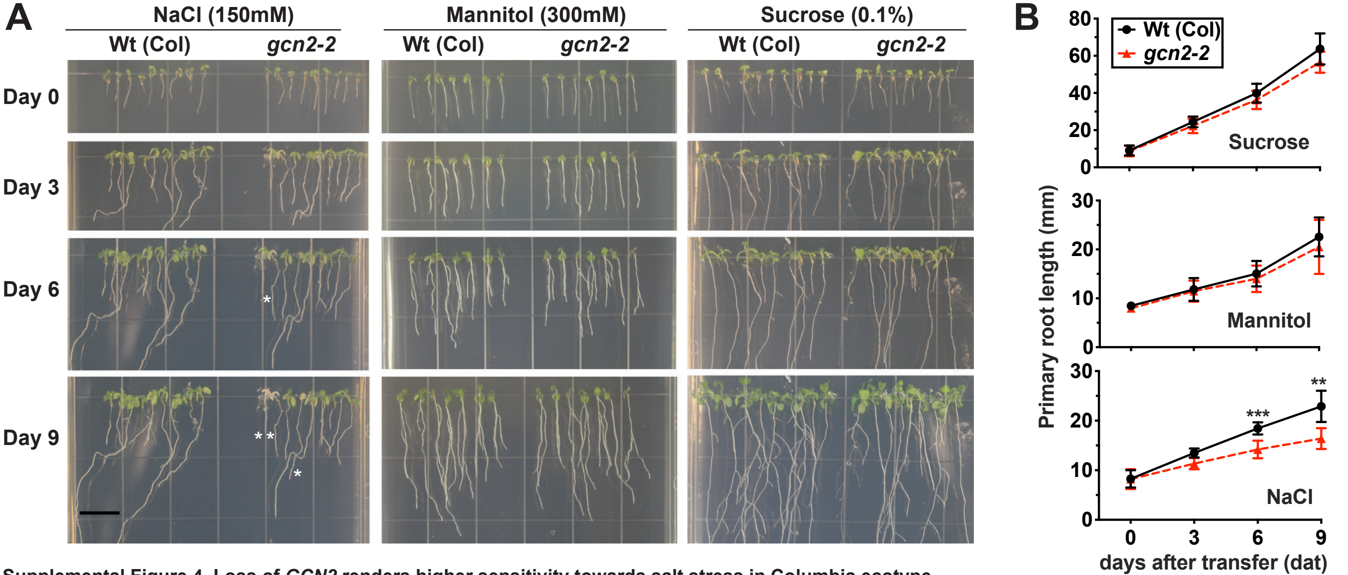


**Supplemental Figure 4. Loss of *GCN2* renders higher sensitivity towards salt stress in Columbia ecotype**

**(A)** Representative images of wild-type Columbia (Wt (Col)) and *gcn2-2* mutant (*gcn2-2*) seedlings grown under 16 hr light and 8 hr dark period on plant media supplemented with 150mM NaCl (salt treatment), 300mM mannitol (osmotic control), or 0.1% sucrose (transfer control). Seedlings were grown on media with 0.1% sucrose for 3 days and transferred to a fresh plate (Day 0). Scale bar is 10mm. Bleached seedlings are marked with an asterisk.

**(B)** Primary root length of Wt and *gcn2-2* mutants from panel (A). Error bars indicate standard error of the mean of four biological replicates with n>36 per experiment (Welch's t-test *P-value <0.05; ** P-value <0.005).
