## Supplemental Figure 5 for "Light dependent activation of the GCN2 kinase under cold and salt stress is mediated by the photosynthetic status of the chloroplast"

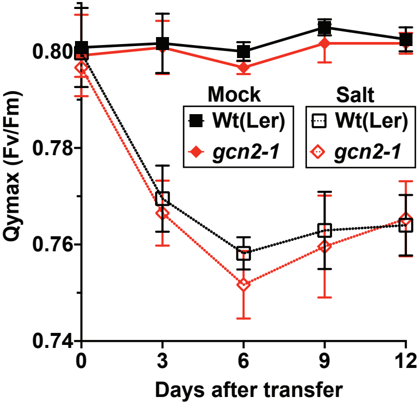


**Supplemental Figure 5 . Effect of salt stress on photosynthetic efficiency of wild-type and *gcn2*.** Time course analysis (Day) of PSII maximal quantum yield (Qymax (Fv/Fm)) of wild-type Landsberg (Wt (Ler)) and *gcn2-1* mutant *(gcn2-1)* seedlings after transfer to 0.1% sucrose (Mock) or 150mM NaCl (Salt) containing media. Error bars represent standard error of the mean from five biological replicates. Welch's t-test *P*-value >0.5.
